# Continued genomic surveillance of *Vibrio cholerae* O1 isolates from cholera cases in Europe, 2023–2024

**DOI:** 10.64898/2026.09.18.752729

**Authors:** Elisabeth Njamkepo, Susann Dupke, Thamida Tauhid, Roger Stephan, Boas van der Putten, Outi Nyholm, Silvia Herrera-León, Richard J. Drew, Martin Cormican, David R. Greig, Borislava Tsafarova, Rosalie Sacheli, Christina Clarke, Ivan N. Ivanov, Claire Jenkins, Maren Lanzl, Inga Fröding, Holger C. Scholz, Caroline Rouard, François-Xavier Weill

**Affiliations:** Institut Pasteur, Université Paris Cité, Unité des Bactéries pathogènes entériques, Centre National de Référence des Vibrions et du choléra, Paris, France; Centre for Biological Threats and Special Pathogens (ZBS 2), National Consultant Laboratory for Human Pathogenic Vibrio species, Robert Koch Institute, Berlin, Germany; Public Health Agency of Sweden, Unit for Laboratory Surveillance of Bacterial Pathogens, Solna, Sweden; Institute for Food Safety and Hygiene, National Reference Center for Enteropathogenic Bacteria and Listeria, Vetsuisse Faculty, University of Zurich, Zurich, Switzerland; Centre for Infectious Disease Control, Institute for Public Health and the Environment (RIVM), Bilthoven, The Netherlands; Microbiology Unit, Finnish Institute for Health and Welfare, Helsinki, Finland; Centro Nacional de Microbiología, Instituto de Salud Carlos III (ISCIII), Madrid, Spain; Rotunda Hospital & Children’s Health Ireland, Dublin, Ireland; School of Medicine, University of Galway, Galway, Ireland; Gastrointestinal Bacteria Reference Unit (GBRU), UK Health Security Agency, London, UK; NRL for Special Bacterial Pathogens, National Center of Infectious and Parasitic Diseases, Sofia, Bulgaria; Clinical Microbiology, National Reference Center for Vibrio spp., University Hospital of Liege, Liege, Belgium; Galway Reference Laboratory Service, University Hospital Galway, Galway, Ireland; NRL for Control and Monitoring of Antimicrobial Resistance, National Center of Infectious and Parasitic Diseases, Sofia, Bulgaria

**Keywords:** Cholera, *Vibrio cholerae*, Europe, 2023, 2024, whole-genome sequencing, microbial genomics, surveillance

## Abstract

**Background:** Following the global resurgence of cholera in 2022, 59 cholera cases were reported by 11 European countries in 2023–2024.

**Aim:** We aimed to confirm that *V. cholerae* O1 isolates associated with these cases belonged to the epidemic seventh pandemic El Tor (7PET) lineage rather than to non-epidemic *V. cholerae* O1 lineages, and to characterise their virulence and antimicrobial resistance (AMR) determinants and phylogenetic relationships.

**Methods:** Fifty *V. cholerae* O1 isolates were available for whole genome sequencing, of which 49 yielded genomes of sufficient quality for analysis. Genomes were analysed together with more than 1,500 publicly available 7PET genomes to place the European isolates into a global phylogenetic context.

**Results:** All 49 genomes belonged to the 7PET lineage and Wave 3. Four sublineages were identified: BD1.2 (2/49), AFR12 (6/49), AFR13 (8/49) and Pre-AFR15 (33/49). The predominance of Pre-AFR15 is consistent with its major contribution to the ongoing global cholera resurgence. In contrast to our 2022 survey, eight AFR13 isolates predicted to be highly drug-resistant were detected, documenting the international spread of this clone carrying genetic determinants associated with resistance to multiple antimicrobial agents recommended for cholera treatment.

**Conclusion:** Whole genome sequencing should be routinely used by reference laboratories to distinguish epidemic 7PET from non-epidemic *V. cholerae* O1 lineages and to monitor the emergence and international spread of clones carrying AMR determinants. Continued collaborative genomic surveillance of travel-associated cholera cases can provide early insights into the emergence, international spread and antimicrobial resistance of 7PET sublineages.

## INTRODUCTION

Cholera is an acute diarrhoeal disease caused by infection with *Vibrio cholerae* O1 and, more rarely, O139 strains carrying the CTXφ prophage encoding cholera toxin [1]. Cholera is transmitted through water or food contaminated with human faeces or by direct faecal–oral transmission. Clinical manifestations range from mild diarrhoea to severe dehydration that can be fatal within hours without prompt oral or intravenous rehydration [1].

The ongoing seventh cholera pandemic began in Indonesia in 1961 and is caused by *V. cholerae* O1 of the El Tor biotype [2]. Since the mid-1960s, South Asia, particularly India and Bangladesh, has been the principal source of repeated international dissemination events, with the pandemic reaching Africa and Europe in 1970 and Latin America in 1991 [3–6]. Following decades of declining incidence, cholera has resurged globally since the early 2020s. According to the World Health Organization (WHO), 472,697 cholera cases from 44 countries were reported in 2022, increasing to 535,321 cases from 45 countries in 2023 and 560,823 cases from 60 countries in 2024 (Figure 1A) [7–9]. During the same period, countries in the WHO European Region reported 52 cholera cases in 2022, 30 cases in 2023 and 29 cases in 2024, almost all associated with recent travel to countries with ongoing cholera transmission (Figure 1B) [7–9].

**Figure 1.**
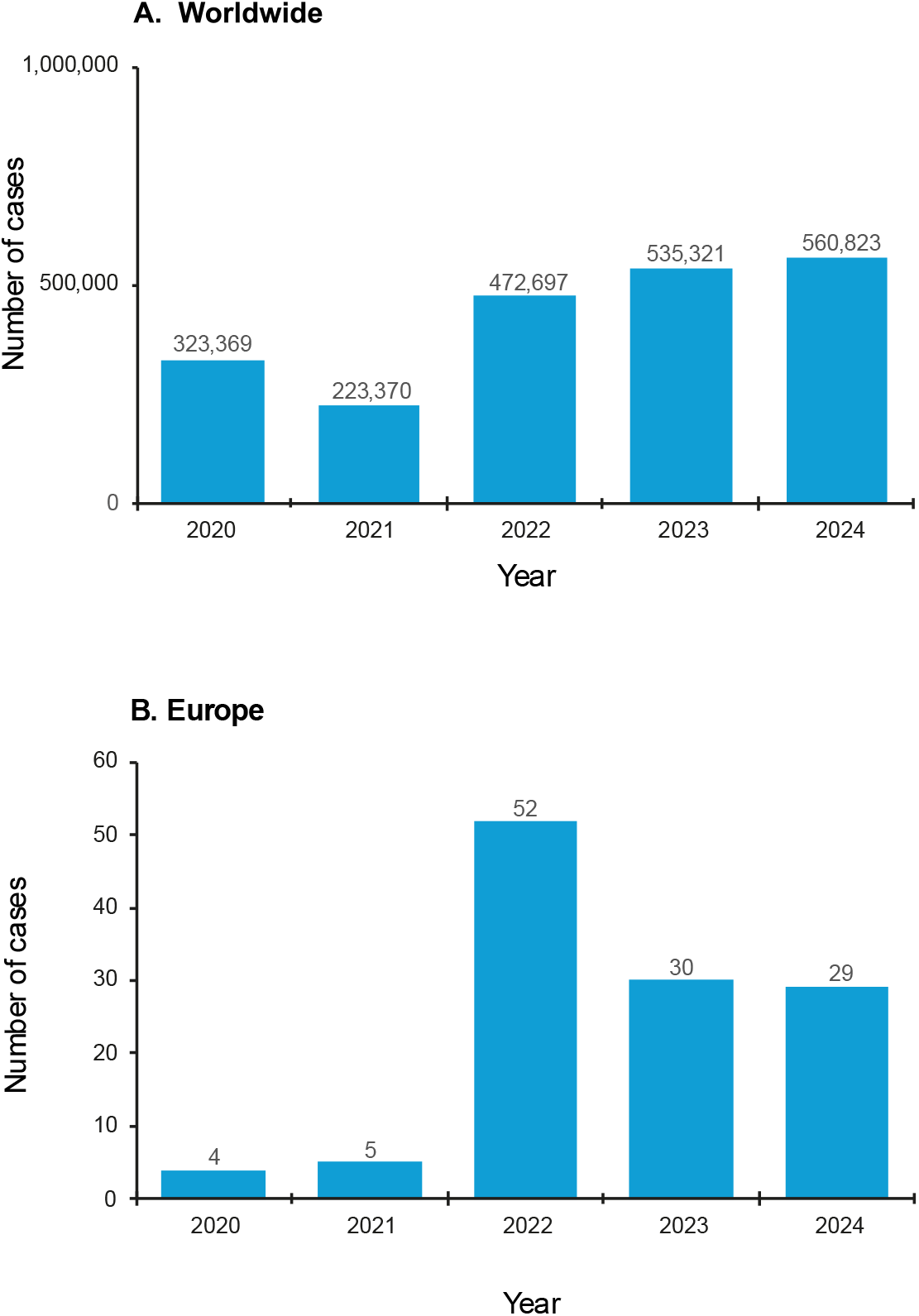
Total number of cholera cases reported to the World Health Organization (WHO) (n = 2,115,580) and cholera cases reported by European countries (n = 120), 2020-2024. Panel A. Countries reporting cholera cases: 2020 (n = 27), 2021 (n = 35), 2022 (n = 44), 2023 (n = 45) and 2024 (n = 60). Panel B. Countries reporting cholera cases: 2020 (n = 2), 2021 (n = 2), 2022 (n = 10), 2023 (n = 6) and 2024 (n = 11, including Andorra). We have retrospectively identified a case of cholera that occurred in Finland in 2022 which had not been mentioned in our previous article (the number of cases in 2022 now stands at 52, spread across 10 reporting countries). Additional information on this isolate (135754) is provided in Supplementary Table 1. In 2023, we have added two cases — one in Belgium and the other in the Netherlands — which had been reported to the ECDC (https://www.ecdc.europa.eu/en/cholera/surveillance-and-disease-data), as well as 14 cases in the UK, to the 14 European cases previously reported to the WHO.

Whole genome sequencing (WGS) has transformed our understanding of the population structure and global dissemination of *V. cholerae* O1. Genomic studies have shown that toxigenic *V. cholerae* O1 strains belonging to the Classical lineage were associated with earlier cholera pandemics [6,10]. In contrast, sustained epidemic transmission during the ongoing seventh cholera pandemic has been associated with toxigenic *V. cholerae* O1 strains belonging to the seventh pandemic El Tor (7PET) lineage [3–6]. Non-toxigenic *V. cholerae* O1 strains and toxigenic strains belonging to non-7PET lineages, including the US Gulf Coast lineage, have been associated with sporadic infections or limited outbreaks, sometimes linked to seafood consumption, with no documented sustained epidemic transmission [6]. Thus, particularly in non-endemic settings, the identification of *V. cholerae* O1, even when toxigenic, does not necessarily indicate that an isolate belongs to the 7PET lineage responsible for the ongoing cholera pandemic and its sustained epidemic transmission. Our previous genomic survey of 49 *V. cholerae* O1 isolates associated with cholera cases reported in Europe in 2022 showed that all belonged to this pandemic lineage [11].

Three successive genomic waves (Waves 1–3) have been described within the 7PET lineage [3], together with numerous phylogenetic sublineages reflecting repeated international dissemination events from South Asia into Africa (sublineages AFR1, AFR3 to AFR15), Europe (EUR1 to EUR8) and Latin America (LAM1 to LAM3) since 1961 [4–6,11]. Between 1970 and 2011, genomic analyses identified at least eight independent introductions of the 7PET lineage into Europe, resulting in outbreaks predominantly in southern and eastern Europe. These introductions originated mainly from Africa for outbreaks in southern Europe and from South Asia for those in eastern Europe, with no evidence of long-term establishment of epidemic 7PET lineages in Europe [5]. WGS also enables the identification and tracking of antimicrobial resistance (AMR) determinants in epidemic lineages [3–6].

In this context, genomic surveillance of *V. cholerae* O1 isolates recovered from cases reported as cholera in Europe provides an opportunity to determine whether these cases are caused by the epidemic 7PET lineage and to monitor the international circulation and AMR of this lineage. Here, we applied WGS to *V. cholerae* O1 isolates recovered from cholera cases reported by countries in the WHO European Region in 2023–2024 to confirm their assignment to the 7PET lineage and to characterise their virulence and AMR determinants. We also investigated their phylogenetic relationships by comparison with more than 1,500 publicly available 7PET genomes. Collaborative genomic surveillance of travel-associated cholera cases provides a sentinel view of the emergence, international dissemination and evolution of epidemic *V. cholerae* lineages, particularly in regions where routine genomic surveillance remains limited.

## METHODS

### Cholera case definition

According to the Global Task Force on Cholera Control (GTFCC), established by WHO, in the absence of an ongoing cholera outbreak—as has generally been the epidemiological situation in European countries since the mid-1990s [5] — a suspected cholera case is defined as a person aged ≥ 2 years presenting with acute watery diarrhoea and severe dehydration, or as a person who died from acute watery diarrhoea with no other known cause of death [12]. A confirmed cholera case is defined as a person with infection by *V. cholerae* O1 or O139 confirmed by culture (including seroagglutination) or by polymerase chain reaction (PCR) for species and serogroup identification [12]. In the absence of an established epidemiological link to another confirmed cholera case or to a known source of exposure, isolates should also be demonstrated to be toxigenic by PCR detection of cholera toxin-encoding genes [12].

### Isolates

Fifty *V. cholerae* O1 isolates were available for genomic analysis from the 59 cholera cases reported by European countries in 2023–2024 [8,9]. These 50 isolates were from the United Kingdom (UK) (n = 23), France (n = 7), Germany (n = 7), Sweden (n = 4), Switzerland (n = 2), the Netherlands (n = 2), Belgium (n = 1), Bulgaria (n = 1), Finland (n = 1), Ireland (n = 1), and Spain (n = 1). Of these 50 isolates, 23 from the UK, one from France and one from Belgium had previously been described in separate reports [13–15].

No *V. cholerae* O1 isolate was available for nine of the 59 cholera cases reported during the study period. These included six of the 13 cases reported by Germany, one of the two cases reported by Spain, and both cases reported by Andorra. In Germany, no culture was performed for one case reported in 2024, and therefore no isolate was available for genomic analysis. The reasons why isolates were unavailable for the remaining five cases, all reported in 2023, could not be determined. In Spain, no further epidemiological or microbiological information was available for the second case, reported from Barcelona in 2024. In Andorra, no bacterial isolates were available from the two cases reported in 2024. These cases had been identified solely by a multiplex PCR assay detecting *V. cholerae,* without targeting the genes encoding cholera toxin. Moreover, neither travel destination (Mexico or the Dominican Republic) was known to have ongoing epidemic cholera transmission at the time, making it unlikely that these cases fulfilled the WHO/GTFCC definition of confirmed cholera.

During the course of this survey, we retrospectively identified one additional *V. cholerae* O1 isolate recovered in Finland in 2022. Because the corresponding case had not been reported to WHO at the time of our previous survey, this isolate (#135754) was not included in that study. Additional information on this isolate is provided in Supplementary Table 1. This isolate was excluded from all analyses restricted to cholera cases reported in 2023–2024. All *V. cholerae* O1 isolates were identified by the participating European reference laboratories according to their respective protocols. Inaba/Ogawa serotyping was performed by slide agglutination for 14 isolates in two of these laboratories.

### Whole-genome sequencing

Whole-genome sequencing was performed by the different participating countries, in accordance with their usual practices, except for the isolate from Belgium, which was sequenced in France. As the genomic sequences initially produced by Sweden were IonTorrent sequences, which was not supported by our bioinformatics pipeline, the four isolates were subsequently resequenced by Novogene (Cambridge, UK) to generate Illumina short-read sequences.

Genomic deoxyribonucleic acid (DNA) was extracted with the Maxwell Cell DNA Purification Kit (Promega, Madison, Wisconsin, USA) (France and Spain), the Maxwell RSC Cultured Cells DNA Kit (France, Sweden and the Netherlands), the QIAamp DNA Mini Kit (Qiagen, Hilden, Germany) (Germany), the DNeasy Blood & Tissue Kit (Qiagen) (Switzerland), the MagAttract HMW DNA Isolation Kit (Qiagen) (Finland), the EZ1&2 DNA Tissue Kit (Qiagen) (Ireland), or the QIAsymphony system (Qiagen) (UK), in accordance with the manufacturers’ recommendations, or using an in-house protocol previously described [16] (Bulgaria). The libraries were prepared with the Nextera XT Kit (Illumina, Inc., San Diego, USA) (France, Finland, Germany and UK), the Illumina DNA Prep Kit (the Netherlands, Spain, Switzerland and Ireland), the Novogene NGS DNA Library Prep Set (Novogene) (Sweden) or the NGS DNA Fragmentation & Library Prep Kit (BioDynami, Huntsville, AL, USA) (Bulgaria). Sequencing was performed with the following Illumina platforms: NextSeq 500 (France), NextSeq 550 (Bulgaria and the Netherlands), NextSeq 1000 (UK and Ireland), NextSeq 2000 (the Netherlands), NovaSeq 6000 (Spain), NovaSeq X Plus Series (Sweden), MiSeq (Finland and Germany), MiniSeq (Switzerland) or HiSeq 2500 (UK), generating 70–150 bp paired-end reads and yielding a mean coverage of 163-fold (minimum, 43-fold; maximum, 507-fold). The list of genomes generated in this study, together with their accession numbers, is provided in Supplementary Table 1.

### Genomic sequence analysis

All reads were filtered with FqCleanER v.23.07 (https://gitlab.pasteur.fr/GIPhy/fqCleanER) to eliminate adaptor sequences and discard low-quality reads with phred scores <28 and a length <70 bp [17]. Taxonomic classification of sequencing reads was performed using Kraken 2 v.2.1.1, and species identification was independently assessed using KmerFinder v.3.0.2 (https://genepi.dk/speciesfinder/) [18]. Genomes were excluded if fewer than 99% of reads were assigned to *V. cholerae* by Kraken 2 and contamination by another bacterial species was detected by KmerFinder. Short reads were assembled with SPAdes v.3.15.5, with default settings in most cases, but with the parameter “--phred-offset 33” for some genomes [19]. Sequence type was determined with the multilocus sequence type scheme of Octavia et al. [20]. The various genetic markers were analysed with the Basic Local Alignment Search Tool (BLAST) v.2.2.26 against reference sequences of the O1 *rfb* gene, *ctxB*, *wbeT*, and the whole locus of VSP-II, as previously described [4]. The presence and type of acquired antibiotic resistance genes (ARGs) or ARG-containing structures were determined with ResFinder v.4.3.3 (https://genepi.food.dtu.dk/resfinder), BLAST analysis against GI-15, Tn*7*, and SXT/R391 integrative and conjugative elements, and PlasmidFinder v.2.1.1 (https://genepi.dk/plasmidfinder). The presence of mutations in the genes encoding resistance to quinolones (*gyrA*, *parC*), resistance to nitrofurans (*nfsA*, nitroreductase, VC0715 and VCA0637, dihydropteridine reductase), or restoring susceptibility to polymyxin B (VC1320, *vprA*) were investigated by manual analysis of the sequences assembled *de novo* with BLAST, as previously described [4,21].

### Additional genomic data

Raw sequence files and assembled genomes from 1,521 7PET strains were downloaded from the European Nucleotide Archive (ENA, https://www.ebi.ac.uk/ena/) or GenBank (https://www.ncbi.nlm.nih.gov/genbank/). The complete list of contextual genomes and their accession numbers is provided in Supplementary Table 2.

### Phylogenetic analyses

The paired-end reads and draft or assembled genomes were mapped onto the reference genome of *V. cholerae* O1 El Tor N16961, also known as A19 (GenBank accession numbers LT907989 and LT907990) with Snippy version 4.6.0/BWA v.0.7.17 (https://github.com/tseemann/snippy). Single-nucleotide variants (SNVs) were called with Snippy v.4.6.0/Freebayes v.1.3.2 (https://github.com/tseemann/snippy), under the following constraints: mapping quality of 60, a minimum base quality of 13, a minimum read coverage of 4, and a 75% read concordance at the locus for a variant to be reported. An alignment of core genome SNVs was produced in Snippy for phylogeny inference. Repetitive (insertion sequences and the TLC-RS1-CTX region) and recombinogenic (VSP-II) regions in the alignment were masked [4]. Putative recombinogenic regions were detected and masked with Gubbins v.3.2.0 [22]. A maximum likelihood (ML) phylogenetic tree was built from an alignment of 11,030 chromosomal SNVs, with RAxML v.8.2.12, under the general time-reversible (GTR) model with 200 bootstraps [23]. This global tree was rooted on the A6 genome — the earliest and most ancestral 7PET isolate, collected in Indonesia in 1957 [4] — and visualised with iTOL v.7.6 (https://itol.embl.de) [24].

## RESULTS

In 2023-2024, 12 countries of the WHO European Region reported 59 cases of cholera (Figure 1), from which 50 *V. cholerae* O1 isolates were available for genomic analysis. Genome sequences for 25 isolates were already available [13–15], and the remaining 25 isolates were sequenced for this study. In total, 24 of the 25 genomes passed quality control (one genome was contaminated with *Escherichia coli* and was removed). The epidemiological metadata (country of infection) and genomic characteristics of the 49 isolates are summarised in Table 1. Additional phenotypic and genomic characteristics of individual isolates are provided in Supplementary Table 1.

**Table 1.** Main characteristics of *Vibrio cholerae* O1 isolates, Europe, 2023-2024 (n = 49)

| Strain name | Isolation year | Reported travel | Sublineage | AMR determinants |
|---|---|---|---|---|
| CNRVC230106 | 2023 | Kenya | AFR13 | *blaPER-7,aadA2,msr(E),mph(A),mph(E),sul1,dfrA1,gyrA_S83I,parC_S85L* |
| CNRVC230266 | 2023 | India | Pre-AFR15 | *strAB,floR,sul2,dfrA1,gyrA_S83I,parC_S85L* |
| CNRVC230274 | 2023 | Kenya | AFR13 | *blaPER-7,aadA2,msr(E),mph(A),mph(E),sul1,dfrA1,gyrA_S83I,parC_S85L* |
| SE-VC2302 | 2023 | Iraq | Pre-AFR15 | *strAB,floR,sul2,dfrA1,gyrA_S83I,parC_S85L* |
| SE-VC2301 | 2023 | Iraq | Pre-AFR15 | *strAB,floR,sul2,dfrA1,gyrA_S83I,parC_S85L* |
| SE-VC2303 | 2023 | Iraq | Pre-AFR15 | *strAB,floR,sul2,dfrA1,gyrA_S83I,parC_S85L* |
| NL23-02 | 2023 | Ethiopia | AFR13 | *blaPER-7,aadA2,msr(E),mph(A),mph(E),sul1,dfrA1,gyrA_S83I,parC_S85L* |
| 1213065 | 2023 | Pakistan | Pre-AFR15 | *strAB,floR,sul2,dfrA1,gyrA_S83I,parC_S85L* |
| 1202169 | 2023 | Pakistan | Pre-AFR15 | *strAB,sul2,dfrA1,gyrA_S83I,parC_S85L* |
| 1113756 | 2023 | Zimbabwe* | AFR13 | *blaPER-7,aadA2,msr(E),mph(A),mph(E),sul1,dfrA1,gyrA_S83I,parC_S85L* |
| 1200093 | 2023 | Unknown | BD1.2 | *strAB,floR,sul2,dfrA1,gyrA_S83I,parC_S85L* |
| 1182832 | 2023 | Pakistan | Pre-AFR15 | *strAB,floR,sul2,dfrA1,gyrA_S83I,parC_S85L* |
| 1182831 | 2023 | Pakistan | Pre-AFR15 | *strAB,floR,sul2,dfrA1,gyrA_S83I,parC_S85L* |
| 1216946 | 2023 | Pakistan | Pre-AFR15 | *strAB,floR,sul2,dfrA1,gyrA_S83I,parC_S85L* |
| 1901530 | 2023 | Kenya | AFR13 | *blaPER-7,aadA2,msr(E),mph(A),mph(E),sul1,dfrA1,gyrA_S83I,parC_S85L* |
| 1241637 | 2023 | Unknown | Pre-AFR15 | *strAB,floR,sul2,dfrA1,gyrA_S83I,parC_S85L* |
| 1232156 | 2023 | Iraq | Pre-AFR15 | *strAB,floR,sul2,dfrA1,gyrA_S83I,parC_S85L* |
| 1232116 | 2023 | Iraq | Pre-AFR15 | *strAB,floR,sul2,dfrA1,gyrA_S83I,parC_S85L* |
| 1260733 | 2023 | Unknown | Pre-AFR15 | *strAB,floR,sul2,dfrA1,gyrA_S83I,parC_S85L* |
| 1282694 | 2023 | Iran | Pre-AFR15 | *strAB,floR,sul2,dfrA1,gyrA_S83I,parC_S85L* |
| 1289734 | 2023 | Pakistan | Pre-AFR15 | *strAB,floR,sul2,dfrA1,gyrA_S83I,parC_S85L* |
| Vib1194 | 2023 | Iraq | Pre-AFR15 | *strAB,floR,sul2,dfrA1,gyrA_S83I,parC_S85L* |
| Vib1193 | 2023 | Pakistan | Pre-AFR15 | *strAB,floR,sul2,dfrA1,gyrA_S83I,parC_S85L* |
| Vib1074 | 2023 | Cameroon | AFR12 | *dfrA1,gyrA_S83I,parC_S85L* |
| Vib1073 | 2023 | Cameroon | AFR12 | *dfrA1,gyrA_S83I,parC_S85L* |
| CNRVC240029 | 2024 | India | Pre-AFR15 | *strAB,floR,sul2,dfrA1,gyrA_S83I,parC_S85L* |
| CNRVC240034 | 2024 | India | Pre-AFR15 | *strAB,floR,sul2,dfrA1,gyrA_S83I,parC_S85L* |
| CNRVC240087 | 2024 | India | Pre-AFR15 | *dfrA1,gyrA_S83I,parC_S85L* |
| CNRVC240249 | 2024 | Pakistan | Pre-AFR15 | *strAB,floR,sul2,dfrA1,gyrA_S83I,parC_S85L* |
| CNRVC240363 | 2024 | India | Pre-AFR15 | *strAB,floR,sul2,dfrA1,gyrA_S83I,parC_S85L* |
| SE-VC24 | 2024 | Iraq | Pre-AFR15 | *strAB,floR,sul2,dfrA1,gyrA_S83I,parC_S85L* |
| NL24-01 | 2024 | Iraq | Pre-AFR15 | *strAB,floR,sul2,dfrA1,gyrA_S83I,parC_S85L* |
| Q30185 | 2024 | Bangladesh | BD1.2 | *strAB,floR,sul2,dfrA1 gyrA_S83I,parC_S85L* |
| N24-3072 | 2024 | India | Pre-AFR15 | *strAB,floR,sul2,dfrA1,gyrA_S83I,parC_S85L* |
| N24-3087 | 2024 | Iraq | Pre-AFR15 | *strAB,floR,sul2,dfrA1,gyrA_S83I,parC_S85L* |
| 1537794 | 2024 | Unknown | Pre-AFR15 | *strAB,floR,sul2,dfrA1,gyrA_S83I,parC_S85L* |
| 1585647 | 2024 | Pakistan | Pre-AFR15 | *strAB,floR,sul2,dfrA1,gyrA_S83I,parC_S85L* |
| 1590742 | 2024 | India | Pre-AFR15 | *strAB,floR,sul2,dfrA1,gyrA_S83I,parC_S85L* |
| 1662639 | 2024 | Ghana | AFR12 | *dfrA1,gyrA_S83I,parC_S85L* |
| 1652134 | 2024 | Nigeria | AFR12 | *dfrA1,gyrA_S83I,parC_S85L* |
| 1614228 | 2024 | Pakistan | Pre-AFR15 | *strAB,floR,sul2,dfrA1,gyrA_S83I,parC_S85L* |
| 1626184 | 2024 | Nigeria | AFR12 | *dfrA1,gyrA_S83I,parC_S85L* |
| 1614229 | 2024 | Unknown | Pre-AFR15 | *strAB,floR,sul2,dfrA1,gyrA_S83I,parC_S85L* |
| BG0524 | 2024 | India | Pre-AFR15 | *strAB,floR,sul2,dfrA1,gyrA_S83I,parC_S85L* |
| 1639745 | 2024 | Nigeria | AFR12 | *dfrA1,gyrA_S83I,parC_S85L* |
| V20240052 | 2024 | Egypt | AFR13 | *blaPER-7,aadA2,msr(E),mph(A),mph(E),sul1,dfrA1,gyrA_S83I,parC_S85L* |
| Vib1242 | 2024 | Tanzania | AFR13 | *blaPER-7,aadA2,msr(E),mph(A),mph(E),sul1,dfrA1,gyrA_S83I,parC_S85L* |
| Vib1270 | 2024 | Eritrea | AFR13 | *blaPER-7,aadA2,msr(E),mph(A),mph(E),sul1,dfrA1,gyrA_S83I,parC_S85L* |
| 141727 | 2024 | Pakistan | Pre-AFR15 | *strAB,sul2,dfrA1,gyrA_S83I,parC_S85L* |

\*a trip to sub-Saharan Africa, probably to Zimbabwe. Additional phenotypic and genomic characteristics of individual isolates are provided in Supplementary Table 1.

### Genomic characterisation and phylogenetic clustering

All 49 isolates belonged to multilocus sequence type ST69, which is characteristic of the 7PET lineage [25], and carried the *ctxB7* allele of the cholera toxin B subunit gene.

To place the European *V. cholerae* O1 isolates in a global phylogenetic context, we analysed them together with over 1,500 publicly available 7PET genomes. The phylogenetic tree was constructed from 11,030 non-recombinant SNVs identified among these 1,571 genomes (49 European isolates from this survey, one European isolate from 2022 that was retrospectively identified, and 1,521 contextual genomes) (Figure 2).

**Figure 2.**
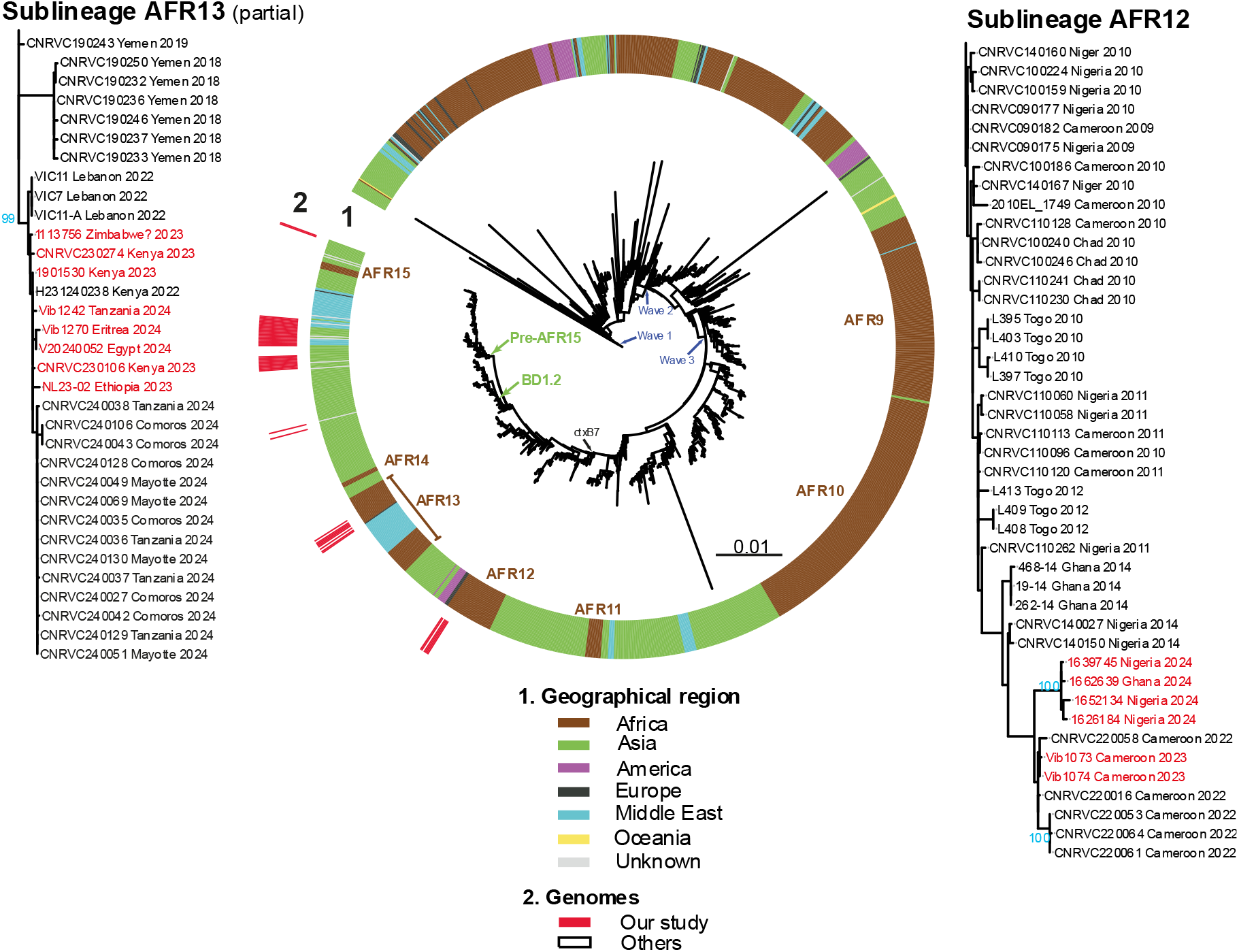
Maximum-likelihood phylogeny of *Vibrio cholerae* O1 El Tor isolates (n = 49), Europe, 2023-2024, compared with 1,522 seventh pandemic *V. cholerae* El Tor genomic sequences (1,521 contextual genomes and one European isolate from 2022). A6 was used as the outgroup. Blue arrows represent the three genomic waves, the black arrow indicates the acquisition of the *ctxB7* allele and green arrows indicate the BD1.2 and Pre-AFR15 sublineages. The colour coding in the inner ring indicates the geographical origins of the isolates, and the most recent African sublineages (AFR10–AFR15) are shown on the right. The 49 isolates studied here are shown in red in the outer ring. A magnification of two selected sublineages is shown on the right (AFR12) and on the left (AFR13, with only the most recent isolates shown), with red text indicating the isolates from 2023–2024 studied. For each genome, the isolate name, country of infection and year of sample collection are indicated at the tree tips. Scale bars indicate the number of nucleotide substitutions per variable site. Selected bootstrap values ≥ 90% are shown in blue.

All 49 isolates belonged to Wave 3 of the 7PET lineage and carried a wild-type *wbeT* (*rfbT*) gene (GenBank accession number JF284685), consistent with the Ogawa serotype [4]. The Ogawa serotype was confirmed by slide agglutination for 14 isolates, whereas serotype assignment for the remaining 35 isolates was based on genomic analysis.

These 49 isolates clustered into four 7PET sublineages: Pre-AFR15 (n = 33), AFR13 (n = 8), AFR12 (n = 6) and BD1.2 (n = 2) (Figure 2).

As in our previous survey, Pre-AFR15 remained by far the predominant sublineage, accounting for 33 of the 49 isolates. These isolates were associated with travel to Pakistan (n = 11), Iraq (n = 9), India (n = 8), and Iran (n = 1). Travel information was unavailable four cases.

Eight isolates belonged to AFR13, a sublineage associated with the emergence and international dissemination of highly drug-resistant isolates. This sublineage was not identified in our previous survey of European cholera cases in 2022 [11]. These isolates were associated with travel to Kenya (n = 3), Tanzania (n = 1), Ethiopia (n = 1), Eritrea (n = 1), Egypt (n = 1), and probably Zimbabwe (n = 1).

Six isolates clustered within AFR12. They were associated with travel to Nigeria (n = 3), Cameroon (n = 2) and Ghana (n = 1). The two isolates associated with travel to Cameroon in 2023 clustered closely with the five Cameroon-associated isolates included in our previous survey (mean pairwise distance: six core-genome SNVs; range 0–13), indicating continued circulation of closely related AFR12 strains in Cameroon. This sublineage has been circulating in West and Central Africa since 2009 [4,26].

Finally, two isolates clustered within the BD1.2 sublineage. One was associated with travel to Bangladesh, whereas no travel information was available for the second isolate, which was recovered in the United Kingdom. This sublineage has been associated with the large cholera outbreak that affected Dhaka in 2022 [27].

### Antimicrobial resistance determinants

All 49 isolates carried mutations in the two chromosomal genes (VC0715 and VCA0637) associated with resistance to nitrofurans and point mutations in the quinolone resistance-determining regions (QRDRs) of the chromosomal genes *gyrA* (S83I) and *parC* (S85L), associated with nalidixic acid resistance and reduced susceptibility to or resistance to ciprofloxacin, as previously described [4,21]. All but six isolates (those belonging to AFR12) had the non-synonymous G265A mutation in *vprA* (VC1320), associated with restored susceptibility to polymyxins [21].

All 49 isolates contained either the complete SXT/R391 genomic element ICE*Vch*Ind5 or a deletion variant thereof [4,28]. The complete ICE*Vch*Ind5 carries genes associated with resistance to streptomycin (*strAB*), sulfonamides (*sul2*), chloramphenicol (*floR*), trimethoprim (*dfrA1*), and trimethoprim–sulfamethoxazole (*sul2* and *dfrA1*). A variant lacking genes ICEVchInd50012–ICEVchInd50015 (including *floR*) (according to GenBank accession number GQ463142) was identified in two Pre-AFR15 isolates. A second variant lacking genes ICEVchInd50011–ICEVchInd50019 (including *strAB*, *floR* and *sul2*) was present in all six AFR12 isolates, all eight AFR13 isolates and one Pre-AFR15 isolate.

Finally, all eight AFR13 isolates harboured an IncC2 plasmid (pCNRVC190243-like; GenBank accession number OW443149.1) carrying genes associated with resistance to aminoglycosides (*aadA2*), sulfonamides (*sul1*), trimethoprim–sulfamethoxazole (simultaneous presence of the plasmid-borne *sul1* gene and the chromosomal *dfrA1* gene), third-generation cephalosporins (extended-spectrum beta-lactamase (ESBL) *bla*_PER-7_), and macrolides (*mph(A)*, *mph(E)*, and *msr(E)*). Thus, all isolates predicted to be highly drug-resistant in this survey belonged to AFR13. For two of these AFR13 isolates (CNRVC230106 and CNRVC230274), previously published phenotypic antimicrobial susceptibility results were consistent with their genomic resistance profiles [13].

## DISCUSSION

Since 2022, the global number of cholera cases reported to WHO has increased, with more than 500,000 cases reported annually in both 2023 and 2024. The number of reporting countries also increased, from 44 in 2022 to 60 in 2024. In contrast, the number of cholera cases reported by European countries decreased from 52 in 2022 to 29–30 in 2023–2024. These figures should nevertheless be interpreted with caution, as they are unlikely to fully reflect the true burden of cholera. Not all countries report cholera cases to WHO. This may be related to limited laboratory capacities, administrative omissions or decisions not to report. Furthermore, under the International Health Regulations, countries are requested to notify WHO of “any event occurring on its territory that may constitute a public health emergency of international concern”; notification of individual sporadic cholera cases is therefore not systematically mandatory [29].

For European cases, additional data sources were required to complement WHO reports, particularly in 2023. While WHO data identified only three European countries reporting a total of 14 cases, information from the European Centre for Disease Prevention and Control (ECDC) Surveillance Atlas of Infectious Diseases (https://www.ecdc.europa.eu/en/cholera/surveillance-and-disease-data) and published literature allowed us to identify three additional countries and 16 additional cases. Since 2024, however, cholera has no longer been included in the ECDC Surveillance Atlas of Infectious Diseases. Imported cholera cases are no longer routinely reported through this system; only events related to domestic transmission are notified to ECDC through EpiPulse (e.g. suspected cholera associated with Ethiopian holy water in 2025). Consequently, our analysis for 2024 was based mainly on WHO reports and informal exchanges with public health microbiologists from reference laboratories across the European Union. These interactions did not identify additional cases. Among the cases reported to WHO in 2024, microbiological confirmation could not be assessed for one patient because the local laboratory did not refer the isolate or sample to the national reference centre. Two other cases reported by a small country without a national reference centre did not meet the WHO/GTFCC criteria for confirmed cholera cases; furthermore, the reported travel histories (Mexico and the Dominican Republic) were not consistent with likely cholera acquisition.

As in our previous survey, the design of this microbiological study and the limited metadata available prevented us from determining whether infected travellers were tourists, business travellers, humanitarian workers, or migrants with historical links between European countries and cholera-endemic regions through colonial or migration pathways.

This study represents the second genomic surveillance investigation based on *V. cholerae* O1 isolates recovered from patients in Europe. Although conducted outside cholera-endemic regions, genomic surveillance of travel-associated cases can provide early warning of the international dissemination of epidemiologically important *V. cholerae* O1 sublineages, particularly antimicrobial-resistant strains and strains associated with large outbreaks. Their value is further increased when genomic surveillance data can be generated rapidly. However, establishing such surveillance remains challenging. In practice, analyses often depend on the annual publication of WHO cholera reports, usually released several months after the end of the surveillance period, followed by additional data collection through reference laboratory networks and review of published literature. The subsequent process of establishing collaborations, obtaining isolate access, sequencing data, or bacterial material through material transfer agreements can further delay analysis. Consequently, genomic surveillance studies may be published several years after the events investigated, reducing their potential impact for real-time public health decision-making.

In our previous survey of cholera cases reported in Europe in 2022 [11], four of the 49 cases for which isolates were available were considered to have been acquired in Europe, all associated with closely related AFR12 strains. In contrast, no locally acquired case was identified among the cases with known travel histories in 2023–2024. However, travel history was unavailable for five of the 50 cases for which isolates were available during this period.

All *V. cholerae* O1 isolates analysed in this study belonged to the 7PET lineage, confirming observations from our previous survey. This finding is important because several other *V. cholerae* O1 lineages exist, including lineages carrying cholera toxin (*ctx*) and toxin-coregulated pilus (*tcp*) genes, but which differ epidemiologically from epidemic cholera by generally causing sporadic infections with limited secondary transmission [6]. One example is the non-7PET *V. cholerae* O1 ST75 lineage, initially described from the US Gulf Coast in the 1970s and subsequently detected in Asia and South Africa [6,30].

In contrast to our previous survey, in which we identified two isolates belonging to Wave 2 (from one case that travelled to Philippines and from one case whose travel information was unavailable), among isolates collected in Europe during 2023–2024, all isolates belonged to the currently circulating third wave of the 7PET lineage.

The Pre-AFR15 sublineage (associated with travel to South Asia and the Middle East) was predominant in our study (33/49) as observed in our previous survey (36/46 high-quality genomes) [11]. The additional Finnish isolate from 2022, identified retrospectively after our previous survey, also belonged to Pre-AFR15. This sublineage has been recognised as a major contributor to the global resurgence of cholera since 2022 and is predominantly associated with transmission in South Asia and the Middle East, with subsequent introductions into eastern and southern Africa [15,31,32]. Interestingly, although Pre-AFR15 accounted for the majority of isolates in our survey, the countries most frequently associated with this sublineage reported relatively few cholera cases. Indeed, 28/33 of the Pre-AFR15 isolates were associated with travel to Pakistan, Iraq or India, although these countries reported at most 1,851 cases annually, whereas Afghanistan reported more than 222,000 suspected cholera cases in 2023 [8,9]. This apparent discrepancy suggests that genomic surveillance of travel-associated cases in Europe may provide complementary information on the geographical distribution of epidemic 7PET sublineages that is not fully captured by national cholera surveillance data, although differences in travel volumes from these countries to Europe and under-ascertainment of cholera in source countries are also likely to contribute.

In this survey, we identified eight AFR13 isolates belonging to the recently described highly drug-resistant clone [33]. Although rehydration remains the cornerstone of cholera treatment, antimicrobial therapy is recommended for severe cases to reduce the duration of diarrhoea, faecal shedding and disease severity [1]. The emergence and international spread of antimicrobial-resistant *V. cholerae* O1 strains therefore represent an increasing public health concern. After its introduction into East Africa around 2013–2014 [21], AFR13 independently acquired IncC2 multidrug-resistance plasmids in Zimbabwe and Yemen (pYA00120881 and pCNRVC190243, respectively) around 2018 [33–35]. Whereas the Zimbabwean clone was not detected again, the Yemeni clone, previously shown to be resistant to two of the three antimicrobial classes recommended for cholera treatment, spread throughout the Middle East and East Africa [13,36–40]. In 2024, this clone caused the cholera outbreak in Mayotte (a French overseas department in the Indian Ocean), involving 261 reported cases [9,13]. In 2025, the same clone was also identified in Europe among patients without recent travel history after consumption of holy water imported from Ethiopia [41–43].

Our eight AFR13 isolates clustered with other isolates belonging to the Yemeni highly drug-resistant clone. Based on their genomic resistance determinants, our AFR13 isolates were predicted to be resistant to two of the three antimicrobial classes currently recommended for cholera treatment, with no known resistance determinants detected for doxycycline. They carried QRDR mutations associated with reduced susceptibility to or resistance to fluoroquinolones and a large IncC2 plasmid harbouring the *mph(A)*, *mph(E)* and *msr(E)* genes, associated with resistance to azithromycin. This IncC2 plasmid also carried the *bla*_PER-7_ gene encoding an ESBL. Six of our AFR13 isolates were associated with travel to East Africa. Three of these isolates, reported independently by three European countries in 2023, contributed to the early evidence that this highly drug-resistant AFR13 sublineage had spread to Kenya [13]. Subsequent genomic analyses of isolates collected during the 2022–2023 cholera outbreak in Kenya documented the local circulation of AFR13 [38,39]. Kenya reported more than 8,800 cholera cases in 2023 [8], highlighting the public health importance of this highly drug-resistant sublineage. Another isolate was associated with travel to Ethiopia, and another with travel to neighbouring Eritrea. Ethiopia reported more than 30,000 cholera cases in 2023 [8]. The detection of AFR13 in a traveller returning from Ethiopia was subsequently supported by genomic data from locally collected Ethiopian isolates, including isolates collected in 2022 [40]. One of our AFR13 isolates recovered in 2024 was associated with travel to Tanzania. Tanzania reported 7,914 cholera cases in 2024 [9], but microbiological data from locally collected outbreak isolates were not available. However, at the beginning of the outbreak in Mayotte in April and May 2024, four imported cases with infections likely acquired in Tanzania yielded highly drug-resistant AFR13 isolates, providing further evidence that this sublineage had spread to Tanzania [13]. The detection of an AFR13 isolate associated with travel to Egypt should be interpreted cautiously, as no cholera cases were reported in the country during that period [8,9]. However, the absence of reported cases does not necessarily exclude undetected or unreported transmission, while travel histories collected during routine surveillance may be incomplete or inaccurate (for example, owing to multiple countries visited during the incubation period or incomplete reporting), and the infection may therefore have been acquired elsewhere. Therefore, the association between the AFR13 isolate and travel to Egypt should be considered uncertain. Taken together, these findings illustrate how genomic surveillance of imported cholera cases in Europe can provide early evidence of the international dissemination of highly drug-resistant lineages and help identify their geographic expansion, particularly in regions where genomic surveillance data are limited.

Our sampling was restricted to isolates from only 12 of the 53 countries covered by the WHO Regional Office for Europe (https://www.who.int/europe/about-us/about-who-europe), representing a relatively small number of cholera cases. Moreover, travel and migration patterns are geographically structured, meaning that travel-associated cases detected in Europe are unlikely to represent a random sample of the global circulation of epidemic *V. cholerae* O1. Consequently, our findings are unlikely to capture the full genetic diversity of 7PET strains circulating worldwide during 2023–2024. Instead, they provide a complementary view of epidemic sublineages sampled through international human mobility and illustrate the value of traveller-based genomic surveillance as an early warning system for emerging epidemic and antimicrobial-resistant clones.

## CONCLUSION

Our findings confirm that travel-associated cholera cases reported in Europe during 2023–2024 were caused exclusively by the epidemic 7PET lineage. In high-income countries where cholera is not endemic, routine laboratory identification of *V. cholerae* O1, even when complemented by detection of cholera toxin genes, cannot reliably distinguish epidemic 7PET strains from non-epidemic toxigenic *V. cholerae* O1 lineages. We therefore recommend that all *V. cholerae* isolates or PCR-positive clinical specimens from suspected cholera cases be referred to national reference laboratories, and that all *V. cholerae* O1 and/or toxigenic *Vibrio* isolates undergo whole-genome sequencing. This would enable discrimination of epidemic 7PET strains, including O1 and O139 serogroups, from non-epidemic *V. cholerae* lineages, while also identifying toxigenic non-O1/non-O139 *V. cholerae* strains, such as O141 [44]. It would also enable the rapid detection of emerging sublineages and antimicrobial-resistant clones. Collaborative genomic surveillance of travel-associated cholera cases in high-income countries therefore represents a valuable sentinel system, providing early warning of the international dissemination and evolution of epidemic 7PET sublineages. Establishing an international genomic surveillance network involving national reference laboratories and public health laboratories from both cholera-endemic and non-endemic countries could further facilitate timely data sharing and provide a broader perspective on the emergence and circulation of epidemic 7PET strains.

## ETHICAL STATEMENT

This study was based exclusively on bacterial isolates and the very limited associated metadata collected by participating reference laboratories under local mandates for the laboratory-based surveillance of cholera in line with local laws and regulations. The associated metadata contained no personal identifiable information and were restricted to the date of isolation and international travel information. Furthermore, the link between the isolates and the participating countries was removed. As a result, neither informed consent nor approval from an ethics committee was required.

## DATA AVAILABILITY

Short-read sequence data were submitted to the ENA (https://www.ebi.ac.uk/ena) or GenBank (https://www.ncbi.nlm.nih.gov/genbank/). The accession numbers of all *V. cholerae* O1 genomes studied are provided in Supplementary Table 2. The whole-genome alignment of the 1,571 genomes and additional files supporting the findings of this study have been deposited in Figshare: https://doi.org/10.6084/m9.figshare.35076607

## ACKNOWLEDGEMENTS

We would like to thank Cecilia Jernberg from the European Centre for Disease Prevention and Control (ECDC) for her support.

## USE OF ARTIFICIAL INTELLIGENCE TOOLS

ChatGPT (OpenAI) was used solely for English-language editing of the manuscript, including grammar, syntax, wording and readability. The authors reviewed and approved all edited text and remain fully responsible for the content, scientific accuracy and conclusions of the manuscript.

## AUTHORS’ CONTRIBUTIONS

F-XW conceptualised and designed the study. EN did the genomic analyses. EN, CR and FXW contributed to data interpretation and visualisation. SD, TT, RS, BvdP, ON, SH-L, RJD, DRG, BT, RS, CC, INI, CJ, MvdB, IF, and HCS were involved in sample collection, metadata curation and genomic sequencing. F-XW drafted the article and all the authors critically reviewed the draft, read and approved the final manuscript.

## COLLABORATORS

Collective authors who fulfil all four ICMJE authorship criteria.

